## Supplementary Figures 1-5 for "Multi-omics analysis of endothelial cells reveals the metabolic diversity that underlies endothelial cell functions"

Supplementary Figure 1

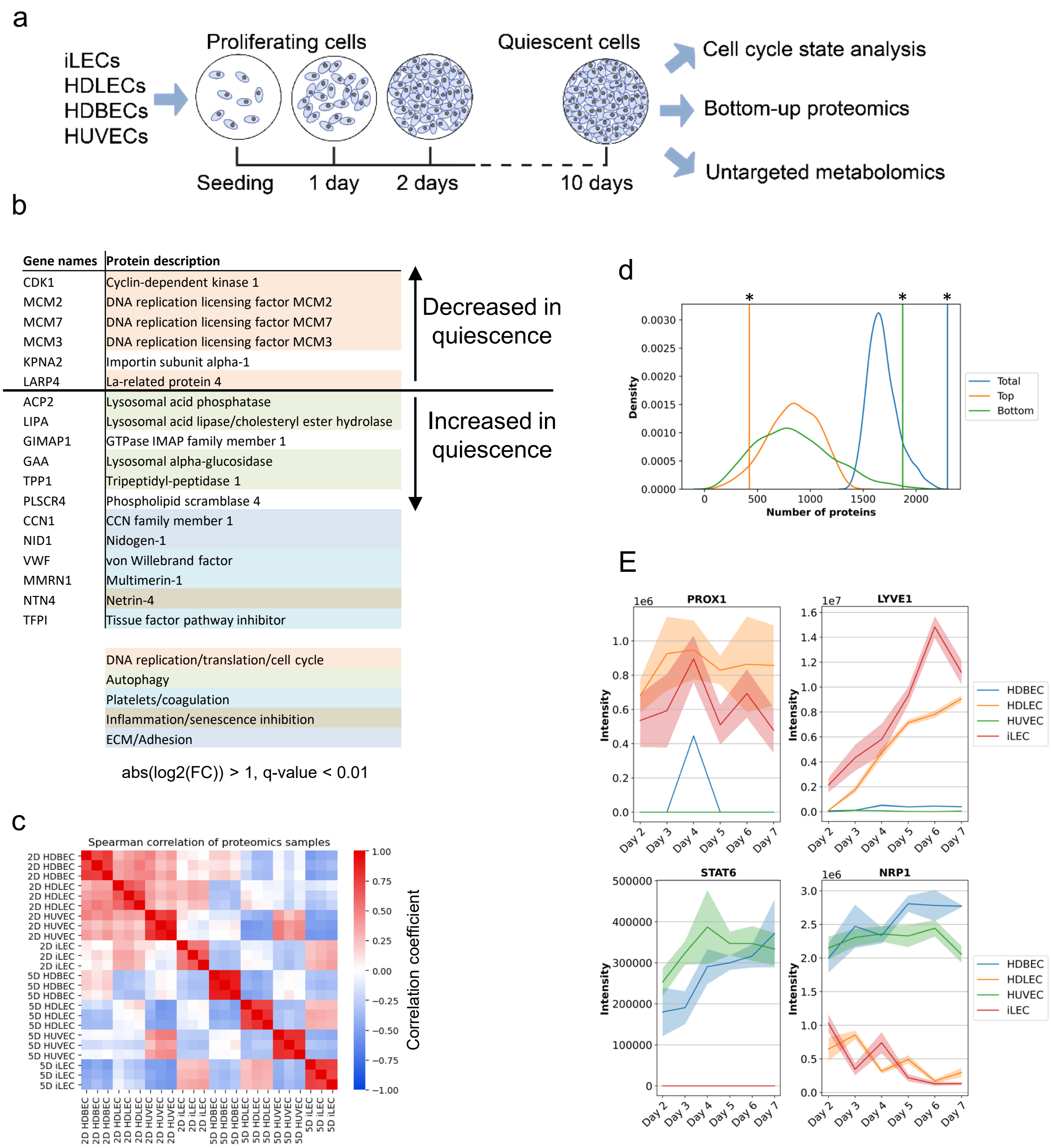


**Supplementary Figure 1 (related to Figure 1).** Proteomic patterns underlying EC identities and states.

(a) Experimental setup. n = 3 replicates per day and cell line for each measurement.

(b) Core protein expression changes. Proteins that pass a threshold of abs(log2(quiescence/proliferation)) > 1 and q-value < 0.01 in all cell lines. The colors indicate the process the proteins are involved in.

(c) Spearman correlation of z-scored proteomics data.

(d) Permutation test to check whether the number of proteins passing the LV1 threshold is random. The curves depict the distributions of the top 10%, bottom 10% and combined numbers. The vertical lines depict the actual values. p-values were determined using a permutation test. * = p-value < 0.01.

(e) Expression levels of LEC markers PROX1 and LYVE1 and BEC markers STAT6 and NRP1.

Supplementary Figure 2

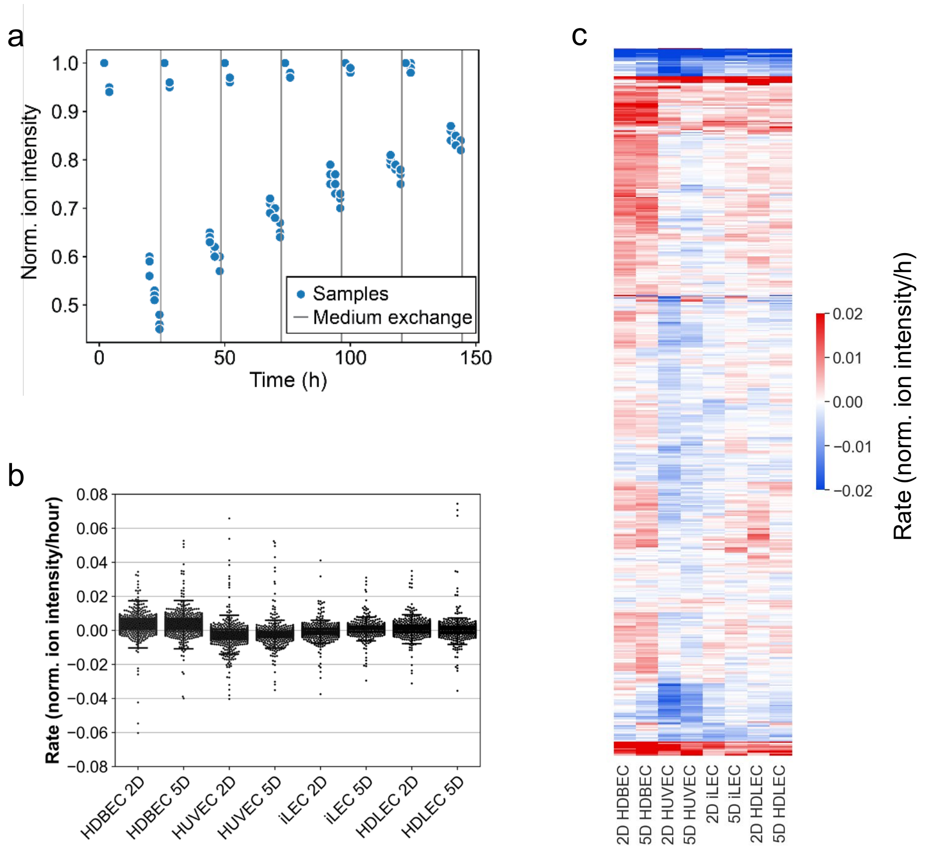


**Supplementary Figure 2 (related to Figure 2).** Overview of analysis and results of extracellular metabolomics data.

(a) Overview of the experimental workflow to determine uptake and secretion rates.

(b) Extend of uptake and secretion rates of all ions in each cell line in proliferation (D2) and quiescence (D5). Negative rates mean uptake, while positive rates mean secretion.

(c) Hierarchical clustered metabolite uptake and secretion rates of all 521 extracellular metabolites measured.

Supplementary Figure 3

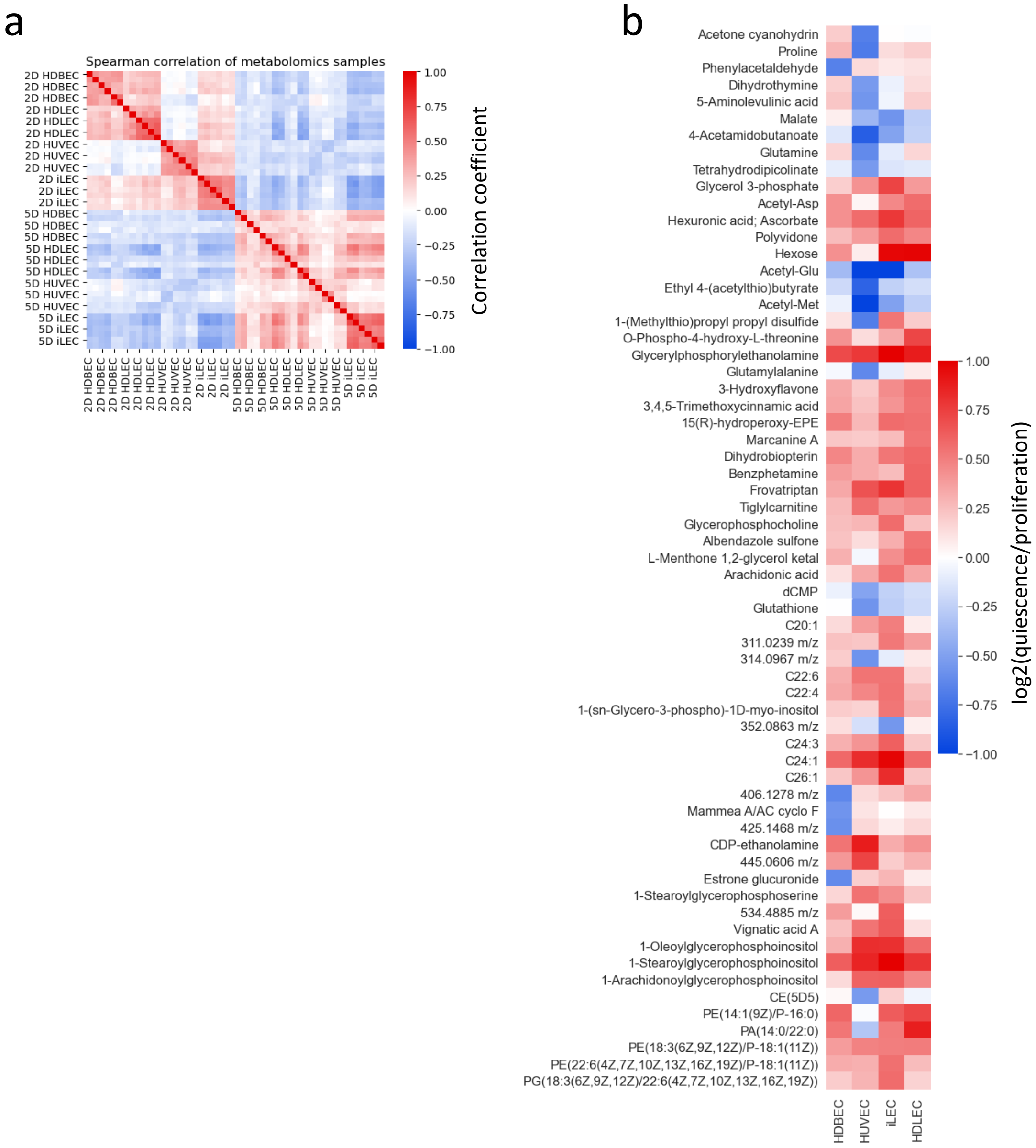


**Supplementary Figure 3 (related to Figure 2).** Intracellular metabolomics patterns.

(a) Spearman correlation of z-scored intracellular metabolomics data.

(b) Overview of metabolites that are changed between quiescence vs proliferation in at least one cell line, passing a threshold of abs(log2(qsc vs prolif)) > 0.5 and adj. p-value < 0.05.

Supplementary Figure 4

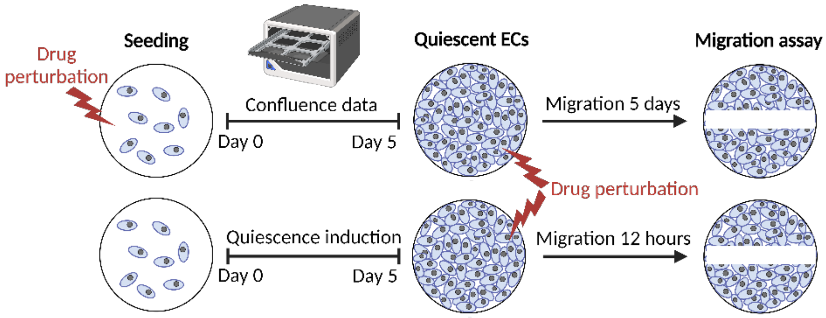


**Supplementary Figure 4 (related to Figure 3).** Overview of the functional validation workflow.

Supplementary Figure 5

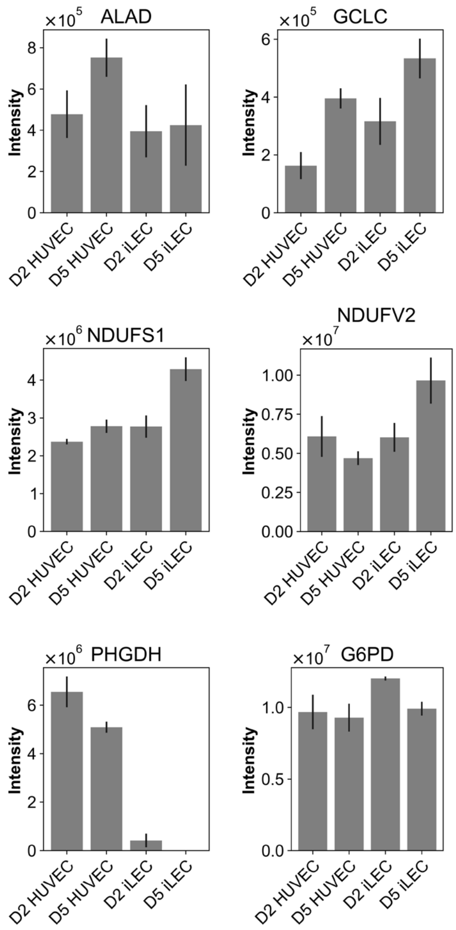


**Supplementary Figure 5 (related to Figure 5).** Expression levels of aminolevulinic acid dehydratase (ALAD, target of SA), glutamate cysteine ligase (GCLC, target of BS), NADH-ubiquinone oxidoreductase 75 kDa subunit (NDUFS1, Complex I, target of rotenone), NADH dehydrogenase [ubiquinone] flavoprotein 2 (NDUFV2, complex I, target of rotenone), D-3-phosphoglycerate dehydrogenase (PHGDH, target of DF) and glucose-6-phosphate 1-dehydrogenase (G6PD, target of G6PDi). Data are represented as mean ± SD.
